# Brief exposure to a diverse range of environmental stress enhances stress tolerance in the polyextremophilic Antarctic midge, *Belgica antarctica*

**DOI:** 10.1101/2020.01.01.887414

**Authors:** J. D. Gantz, B. N. Philip, N. M. Teets, Y. Kawarasaki, L. J. Potts, D. E. Spacht, J. B. Benoit, D. L. Denlinger, R. E Lee

**Affiliations:** Department of Biology and Health Sciences, Hendrix College, Conway, AR USA; Department of Biology, Miami University, Oxford, OH 45056, USA; Department of Entomology, University of Kentucky, Lexington, KY USA; Department of Biology, Gustavus Adolphus College, Saint Peter, MN USA; Department of Evolution, Ecology, and Organismal Biology, The Ohio State University, Columbus, OH 43210, USA; Department of Biological Sciences, University of Cincinnati, Cincinnati, OH, USA

## Abstract

Insects use rapid acclimation to enhance their tolerance of abiotic stresses within minutes to hours. These responses are critical adaptations for insects and other small ectotherms to tolerate drastic changes in temperature, hydration, or other factors that can fluctuate precipitously with ambient conditions or as a result of behavior. Rapid cold-hardening, where insects use brief exposure to modest chilling as a cue to enhance their cold tolerance, is the most thoroughly-studied of these responses and relatively little is known about rapid acclimation that is either triggered by or enhances tolerance of other abiotic stresses. Here, we used larvae of the Antarctic midge, *Belgica antarctica*, a polar extremophile that routinely experiences numerous stresses in nature, to investigate how 2 h exposure to modest environmental stresses affect stress tolerance in insects. Brief pretreatment by various stresses, including hyperosmotic challenge, hypoosmotic challenge, acidity, basicity, and UV irradiation enhanced stress tolerance in *B. antarctica* larvae relative to untreated controls. These results indicate that numerous environmental cues can trigger rapid acclimation in insects and that these responses can enhance tolerance of multiple stresses.

## Introduction

Insects can enhance their stress tolerance within minutes in a process called rapid cold-hardening (RCH) (Lee and Denlinger, 2010). Though chilling is the most commonly-used signal to induce RCH, we have come to understand that these responses are triggered by multiple environmental cues and can exert effects independently of temperature change (Coulson and Bale, 1991; Yoder et al., 2006; Levis et al., 2012). Indeed, these rapid acclimatory responses can also be triggered by modest changes in humidity, and by hypoxia (Coulson and Bale, 1991; Levis et al., 2012). They protect insects from a variety of insults by enhancing tolerance of high temperature, low temperature, freezing, and anoxia, while also increasing resistance to dehydration (Lee et al. 1987; Chen et al., 1991; Coulson and Bale, 1991, Yoder et al., 2006; Lee et al., 2006; Gantz et al., *in review*). The protection afforded by RCH manifests in numerous ways, including reducing cell death, increasing organismal survival, and preserving normal reflexes and behaviors (Lee et al., 1987; Kelty et al., 1996; Kelty and Lee, 1999; Yi and Lee, 2003; Shreve et al., 2004).

In nature, insects can experience abrupt changes in temperature, hydration state, and oxygen availability (Stevenson, 1985; Gringorten and Friend, 1982; Heinrich, 1993; Hoback et al., 1998), suggesting that rapid acclimation is important to overcome the challenges associated with these fluctuations. Even modest decreases in temperature can render insects incapable of flight, courtship, and mating, and RCH helps to mitigate these negative effects (Taylor, 1963; Heinrich, 1972; Shreve et al., 2004). In addition to temperature, dehydration, and anoxia, there are many abiotic stresses that have not yet been investigated as potential triggers for RCH. Natural habitats can present a variety of stresses, including osmotic extremes (Herbst et al., 1988; Nicholson, 1998; Benoit and Denlinger, 2010), acidic and alkaline environments (Winterbourn, 1998; Aislabie et al., 2004; Spitzer and Danks, 2006; Slessarev et al., 2016), starvation and nutrient deprivation (Scott et al., 2004; Simpson et al., 2006), and UV irradiation (Mazza et al., 1999). Since arthropods have successfully colonized nearly all terrestrial habitats on Earth, it is likely that many species are frequently exposed to these less-studied abiotic stresses. In addition to temporal variation, environmental conditions vary in space, suggesting that highly mobile species may need rapid acclamatory processes as they move in and out of certain habitats.

The Antarctic midge, *Belgica antarctica*, lives on the western Antarctic Peninsula in environments where they may be exposed to multiple abiotic stresses at any time. Because of their small size and relative immobility, these midges have few options to deal with environmental stresses other than to tolerate them (Teets and Denlinger, 2014; Lee and Denlinger, 2015). Though temperatures are relatively mild, freezing remains a possibility any day of the year (Lee and Denlinger, 2015). Hydric stress is an even greater challenge for terrestrial arthropods in Antarctica (Kennedy, 1993; Worland and Block, 2003). During the austral summer, fresh water from rain and snowmelt forms pools that inundate many habitats, leading to hypoosmotic conditions and overhydration (Baust and Lee, 1982). Midges also face dehydration and hyperosmotic stress throughout the year (Baust and Lee, 1987; Elnitsky et al., 2009). Water is frozen and thus biologically-unavailable during the non-summer months, as temperatures are constantly below zero. Additionally, meltwater pools can become hyperosmotic as they receive effluent from nearby penguin rookeries and seal wallows; the effects of which are exacerbated by evaporation, which increases solute concentrations as water is lost (Lee and Denlinger, 2015). This same detrital effluent can also cause drastic changes in pH, whereby midges have been found in pools with a pH as low as 4 (Baust and Lee, 1983).

*B. antarctica* is a polyextremophile that tolerates a wide-range of environmental stresses in nature, including freezing to at least -13°C at any time of the year, dehydration to 35% of their original mass, and prolonged exposure to anoxia, hypoosmotic conditions (i.e., immersion in fresh water), hyperosmotic conditions (i.e., immersion in 0.5 M NaCl), and pH exposures between 3 and 12 (Lee and Denlinger, 2015). Consequently, we assessed whether diverse abiotic stresses trigger rapid acclimation by measuring survival of injurious freezing and other stresses after brief exposure to the following cues: starvation, chilling and freezing at modest sub-zero temperatures, high temperature, dehydration, hypoosmotic stress in ultrapure water, hyperosmotic stress in seawater, low pH, high pH, and ultraviolet irradiation.

## Methods

### General notes

This study is made up of two independent investigations that were not originally designed to be used together. As such, treatment conditions differ among assays according to their original group. The assays in this study investigate two primary ideas, 1) whether brief exposure to various abiotic stresses causes rapidly enhanced freezing tolerance, and 2) whether brief exposure to various abiotic stresses causes rapidly enhanced tolerance of numerous other environmental stresses. Assays investigating the effects of various pre-treatments on freezing tolerance, herein called freezing assays, used 2 h exposure to -15°C for as the discriminating cold exposure, and survival was assessed after 2 h of recovery at 2°C. In contrast, assays investigating whether various stresses enhance tolerance to other abiotic stresses, herein called generalized rapid acclimation assays, were exposed to numerous discriminating stress exposures for 24 h and survival was assessed after 24 h of recovery at 2°C. Freezing assays used 30 larvae for assessment of organismal survival and 12 for assessment of cellular damage for each treatment, while generalized rapid acclimation assays used 60 larvae per treatment and measured organismal survival only.

### Insect collection

*B. antarctica* larvae were collected from Torgersen Island, near Palmer Station, Antarctica (64° 46’ S, 64° 04’W) in December 2017. They were maintained in their native substrate at 2°C and constant light for a minimum of 1 week before experimentation. Fourth instar larvae were separated from the substrate by hand immediately before exposure to pretreatment conditions.

### Experimental treatments: freezing protocol

For discriminating cold exposures, larvae were taken directly from their substrate and placed in 1.7 ml microcentrifuge tubes, with 10-12 larvae per tube, 200 µl of water, and a small piece of ice to inoculate ice formation near their freezing point (Lee et al., 2006). Freezing conditions, which varied between assays as described above, were achieved using a programmable cold bath. This injurious freezing protocol was applied to all pretreatment groups immediately after the respective pretreatments concluded. Conditions for freezing assays were 2 h exposure to -15°C followed by 2 h of recovery at 2°C, while generalized rapid acclimation assays used 24 h exposure to -14°C followed by 24 h recovery at 2°C.

### Experimental treatments: food deprivation, temperature, and dehydration

Since larvae were deprived of food during all treatment conditions, we also assessed survival in a second control condition without access to food without exposure to additional treatment conditions in freezing assays. Larvae in this nutrient deprivation control were maintained on damp filter paper in a sealed, plastic petri dish at 4°C. In this control and all temperature and dehydration treatments in both the freezing and the generalized rapid acclimation assays, larvae were exposed to pretreatment conditions for 2 h in groups of 10-12 individuals. For RCH in a supercooled state, larvae were exposed to -5°C in a dry 1.7 ml microcentrifuge tube below their freezing point in the absence of ice formation (Teets et al., 2011), while for RCH in a frozen state, larvae were exposed to -5°C with water and a small piece of ice to inoculate freezing. Larvae exposed to high temperature pretreatments were placed in a 1.7 ml microcentrifuge tube on damp filter paper and placed in a programmable cold bath at either 15, 25, or 30°C. Larvae in dehydration treatments were placed in 1.7 ml tubes with fine, nylon mesh in place of the lid and maintained in sealed desiccating chambers at either 85 or 35% RH, using saturated salt solutions of KCl (85% RH) or MgCl2 (33% RH) (Rockland, 1960).

### Experimental treatments: iso-osmotic, hypoosmotic, hyperosmotic, and pH

Larvae in the iso-osmotic control condition were immersed in Coast’s solution (Coast, 1988) that was diluted to match the osmotic pressure of *B. antarctica* hemolymph at 400 mOsm kg^-1^ (Elnitsky et al., 2009). Hypoosmotic treatments were immersed in ultrapure water (∼0 mOsm kg^-1^), and hyperosmotic treatments were immersed in seawater (∼1000 mOsm kg^-1^) or solutions of NaCl at concentrations of 1.5, 3, and 4 M. Low and high pH solutions were created by titrating Coast’s solution to a pH of 3 and 12 using HCl and NaOH, and diluting the resulting solution to 400 mOsm kg^-1^. Iso-osmotic controls and all pretreatments lasted 2 h.

### Experimental treatments: UV

Larvae in UV treatments were maintained in glass jars with damp filter paper and placed outside in direct sunlight. Jars were submerged up to their openings in an ice bath to control for temperature. The jar used for the UV control treatment was wrapped in duct tape and covered with an opaque lid. The experimental treatment did not receive an opaque wrapping and, instead of a lid, the opening was covered with fine nylon mesh. Treatment lasted 4 h. Temperature was measured in each jar immediately before and after treatment using a copper-constantin (type J) thermocouple and a hand-held reader. At both time points, temperatures in each jar were between 0.0 and 0.5°C.

### Assessing survival

Organismal survival was assessed either 2 or 24 h after removal from the discriminating stress exposure. Larvae were scored as alive if they could be stimulated to move by touch. Cell survival was assessed by adapting a LIVE/DEAD sperm viability kit (Molecular Probes, Eugene, OR, USA) to discriminate between undamaged and freezing-damaged cells (Yi and Lee, 2003). Briefly, excised fat body and midgut tissues were incubated in a membrane-penetrating green nuclear stain (SYBR-14) and an impermeant red nuclear stain (propidium iodide), and numbers of red and green cells were counted via fluorescent microscopy. Cells with membranes that were compromised by freezing damage fluoresced red and were scored as dead, while cells with intact membranes fluoresced green and were scored as alive. We then scored a minimum of 100 cells for each tissue type, from 12 larvae, and calculated the ratio of live vs. dead cells.

### Statistical analyses

For freezing assays, organismal survival data were fitted to general linear models (three models, one for each respective control), each with a logit link function and a binomial error distribution. We followed each with Tukey’s HSD to control for pairwise comparisons (R Foundation for Statistical Computing, Vienna, Austria). However, we only report significance (α = 0.05) for Tukey-corrected pairwise comparisons to our three controls, respectively. Cell survival data were analyzed using a one-way analysis of variance (ANOVA) on the mean proportion of undamaged cells per larva. The Holm-Sidak method was used for post-hoc pairwise comparisons, and data are reported as mean ± SEM. Cellular survival distributions that failed the Shapiro-Wilk test of normality were analyzed using Kruskal-Wallis one-way ANOVA on ranks with post hoc Tukey test for pairwise comparisons and are reported as medians with their interquartile range (IQR).

For generalized rapid acclimation assays, data were fitted to general linear models and subsequent planned contrasts with Bonferroni correction.

## Results

### Organismal survival: freezing assays

When only exposed to the injurious freezing treatment, the survival was 0.1, which was not significantly different compared to starvation for 2 h, at 0.2 (Fig. 1A). Pretreatment with RCH supercooled, RCH frozen, moderate dehydration, and severe dehydration enhanced organismal survival relative to starved controls, at 0.83, 0.80, 0.70, and 0.73 respectively (*p* < 0.01; Fig. 1A). While survival in iso-osmotic control larvae was 0.17, immersion in hypoosmotic and hyperosmotic solutions increased the rate to 0.63 and 0.83 (*p* < 0.01; Fig 1B). Low and high pH solutions did not significantly increase the rate of organismal survival, at 0.33 and 0.5, respectively (Fig. 1B). Finally, larvae that experienced UV irradiation survived at a rate of 0.33 compared to 0.03 in the UV controls (*p* < 0.05; Fig. 1C).

**Figure 1.**
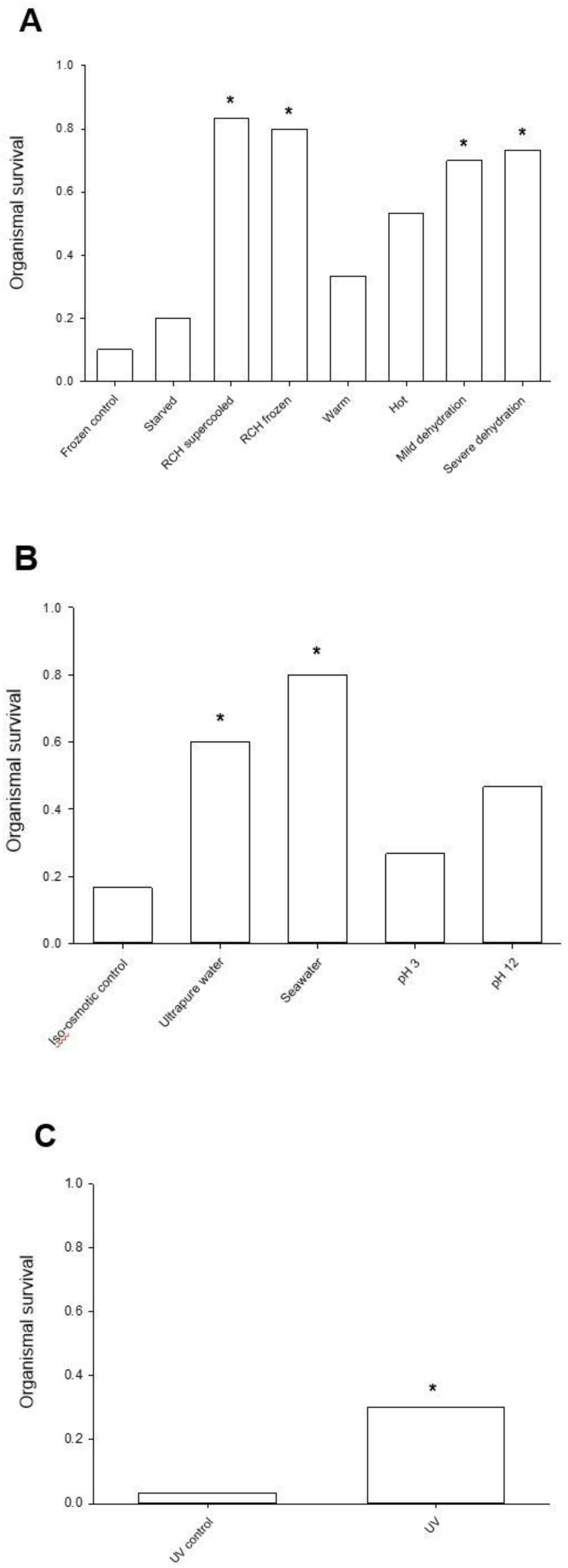
Brief exposure to diverse abiotic stresses enhanced organismal freeze tolerance. **A**. Low temperature and dehydration cues, but not high temperature, enhanced freeze tolerance relative to starved control. Frozen controls are shown for reference, though were not significantly different from starved. * denotes significance relative to starved treatment only (*p* < 0.01). **B**. Exposure to osmotic perturbation in ultrapure water and seawater enhanced freezing relative to iso-osmotic control (*p* < 0.01). * denotes significance relative to iso-osmotic control. **C**. Exposure to UV irradiation enhanced freeze tolerance relative to UV control (*p* < 0.05).

### Cell survival: starvation, temperature, and dehydration

Starvation for 2 h enhanced the rate of survival in both midgut and fat body tissue, at 0.28 (IQR = 0.1) and 0.22 ± 0.04 compared to control treatments, at 0.19 (IQR = 0.1) and 0.08 ± 0.03, respectively (*p* < 0.05; Fig. 2). Since the other treatments within this group were starved while exposed to pretreatment conditions, we compared the rates of survival for each treatment to the starved group rather than to those that were only frozen at -15°C without pretreatment. Midgut and fat body tissues exhibited similar trends among all of these treatments (RCH supercooled, RCH frozen, moderate high temperature, extreme high temperature, moderate dehydration, and severe dehydration; Figs. 3 and 4). Moderate high temperature was the only treatment that did not enhance freeze tolerance in either tissue type, and severe dehydration was the only treatment that increased survival in one tissue (midgut) and not in the other (fat body). Each of the remaining treatments enhanced freeze tolerance relative to starved controls, with 2-3-fold increases in the rates of survival of RCH supercooled, RCH frozen, and mild dehydration in both tissue types (*p* < 0.001).

**Figure 2.**
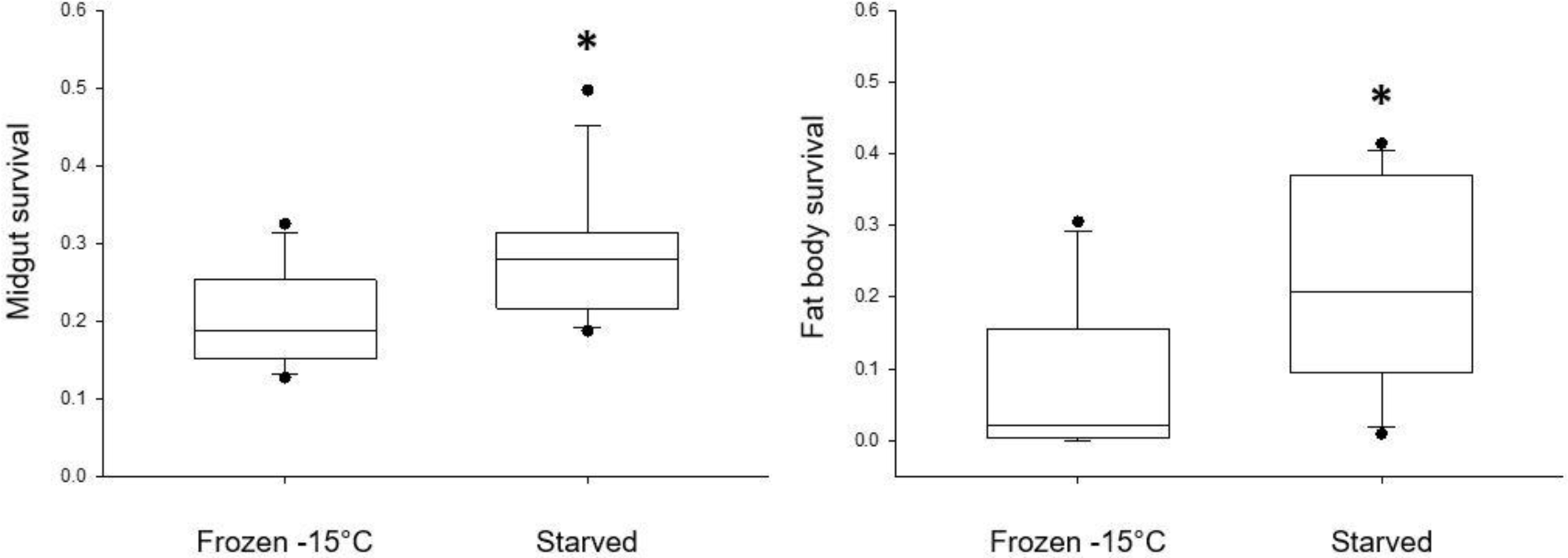
Starvation reduced cellular damage. Ratio of surviving midgut and fat body tissues, relative to frozen control. * denotes significance at *p* < 0.05.

**Figure 3.**
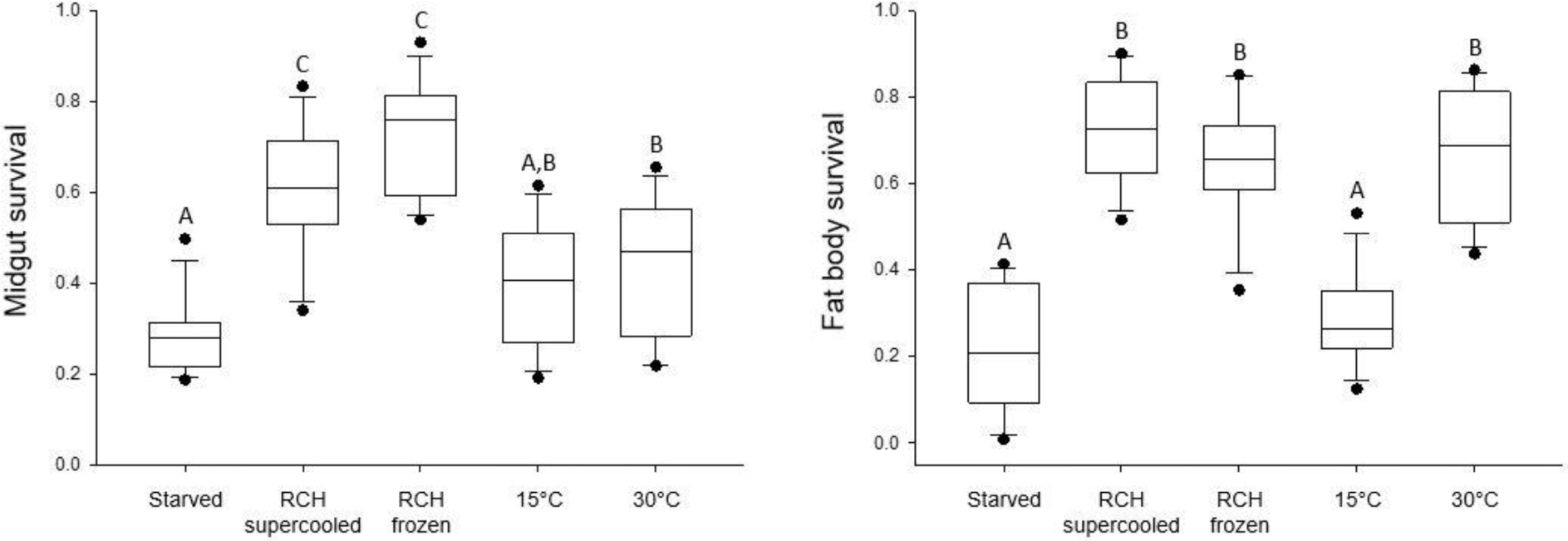
High and low temperature perturbation reduced cellular damage in midgut and fat body cells. Ratio of surviving midgut and fat body tissues relative to starved control. Treatments not sharing a letter were significantly different (*p*<0.05).

**Figure 4.**
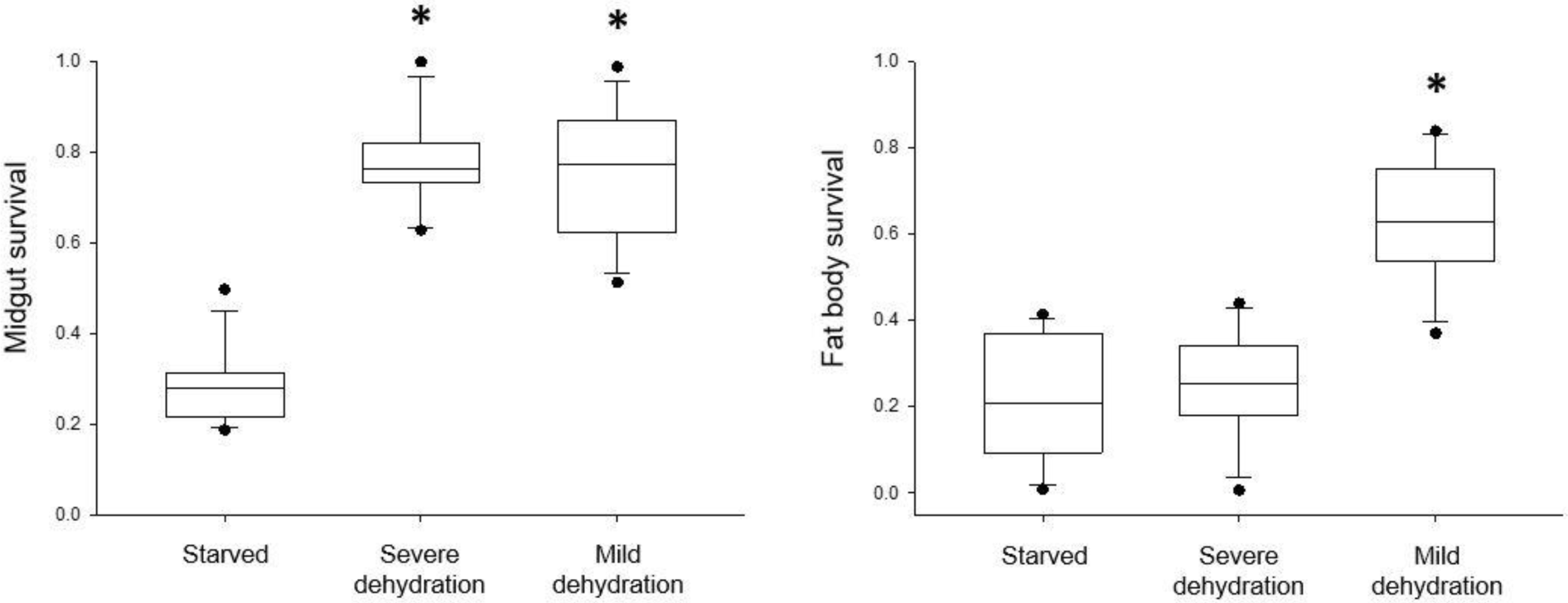
Dehydration reduced cellular damage in midgut and fat body tissues. Ratio of surviving midgut and fat body tissues, relative to starved control. * denotes significance at *p* < 0.01.

### Cellular survival: hypoosmotic, hyperosmotic, and pH

Midgut and fat body cells in iso-osmotic control treatments survived freezing at rates of 0.32 ± 0.03 and 0.12 ± 0.03, respectively. In midgut tissue, immersion in ultrapure water (hypoosmotic) and seawater (hyperosmotic) caused 2.25- and 1.8-fold increases in survival (*p* < 0.001; Fig. 5). The effects of these treatments were even larger in fat body tissue, where ultrapure water and seawater produced 4.67- and 4.25-fold increases in the rate of survival, respectively (*p* < 0.001; Fig. 5)). Immersion in an acidic solution caused a 1.6-fold increase in survival in midgut tissue (*p* < 0.05), though this had no effect on fat body survival (Fig. 6). Immersion in an alkaline solution had no effect on the survival rate of midgut tissue while producing a 4.2-fold increase in fat body survival (*p* < 0.001; Fig. 6).

**Figure 5.**
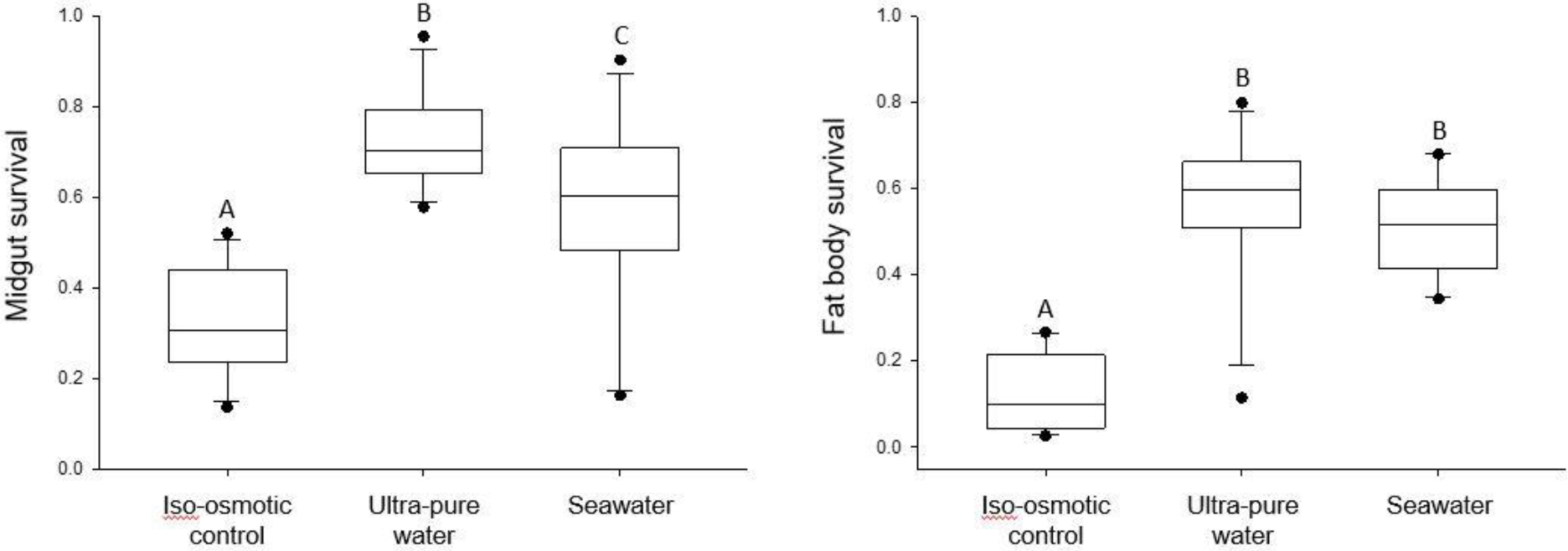
Osmotic perturbation in ultrapure water and seawater reduced cellular damage in midgut and fat body tissues. Ratio of surviving midgut and fat body cells relative to iso-osmotic control. Treatments not sharing a letter were significantly different (*p*<0.05).

**Figure 6.**
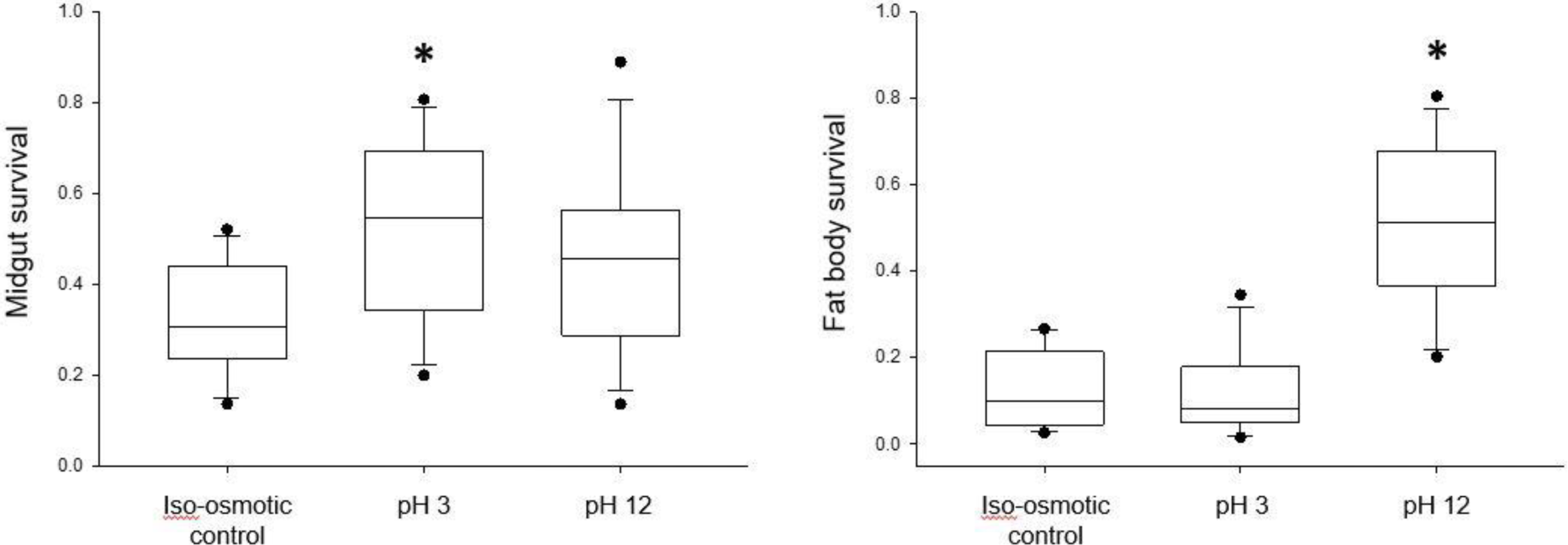
Exposure to high and low pH reduced cellular damage in midgut and fat body tissues. Ratio of surviving midgut and fat body cells relative to iso-osmotic control. * denotes significance at *p* < 0.05.

### Cellular survival: UV

Cellular survival in midgut and fat body tissues in the UV control treatment were 0.63 ± 0.06 and 0.09 ± 0.04, respectively. UV exposure had no effect on survival in midgut tissue, but caused a 4.1-fold increase in survival rate of fat body cells (0.37 ± 0.02; *p* < 0.001; Fig. 7).

**Figure 7.**
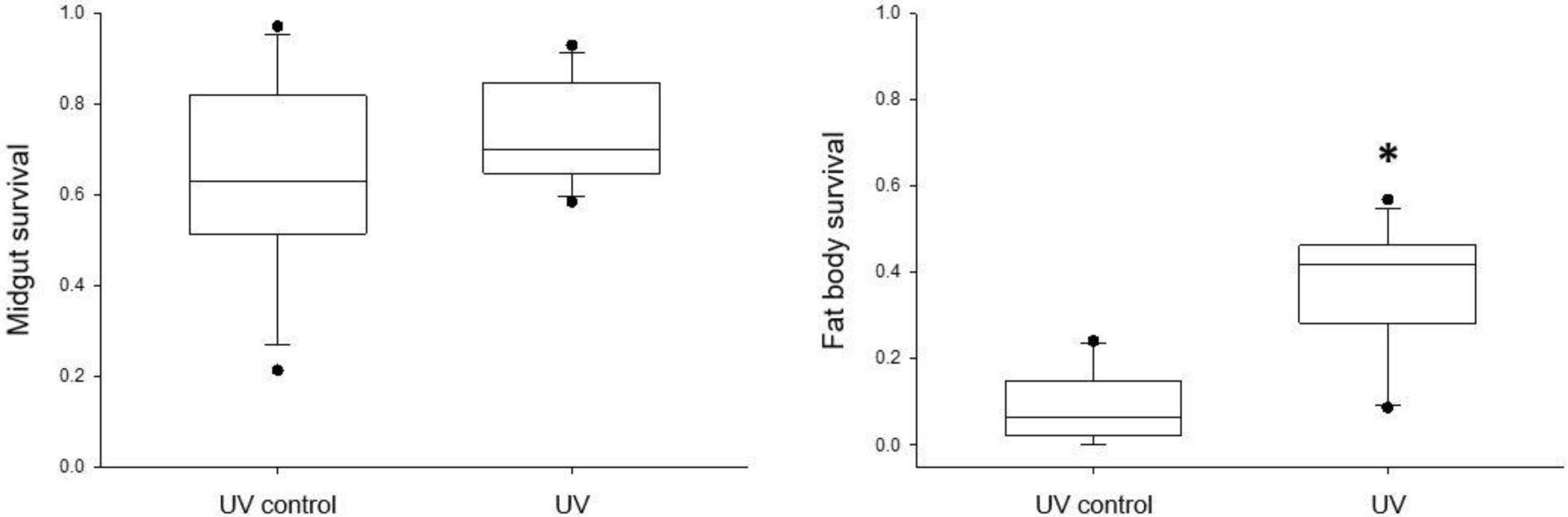
Exposure to solar radiation reduced cellular damage in fat body tissue. Ratio of surviving midgut and fat body cells relative to UV control. * denotes significance at *p* < 0.05.

### Generalized rapid acclimation assays

These assays used a factorial design to investigate organismal survival of four discriminating stress exposures (freezing at -14°C, high temperature at 30°C, desiccation at 35% RH, and hyperosmotic salinity at 4 M NaCl) following 2 h pretreatment by six modest, abiotic stresses (freezing at -5°C, high temperature at 25°C, dehydration at 85% RH, osmotic perturbation immersed in either hypoosmotic water near 0 mOsm kg^-1^ or in a hyperosmotic, 1.5 M NaCl solution, and acid at pH 3). In larvae exposed to discriminating cold, pretreatments of high temperature, hypoosmotic challenge, and acidity resulted in survival at 0.72, 0.66, and 0.57, respectively, which were significantly greater than untreated controls at 0.28. Following discriminating heat exposures, pretreatment with high temperature, 0.87, and desiccation, 0.7, enhanced survival relative to controls, 0.38. In contrast, freezing, hypoosmotic challenge, hyperosmotic challenge, and acidity all decreased survival relative to controls, at 0.07, 0.0, 0.05, and 0.1. In larvae exposed to discriminating desiccation, pretreatment of dehydration, hypoosmotic challenge, and hyperosmotic challenge resulted in survival at 0.57, 0.65, and 0.5, respectively, each of which was significant relative to survival in controls at 0.25. Larvae in the discriminating hyperosmotic environment demonstrated no differences among between treatments and the control (Fig. 8).

**Figure 1.**
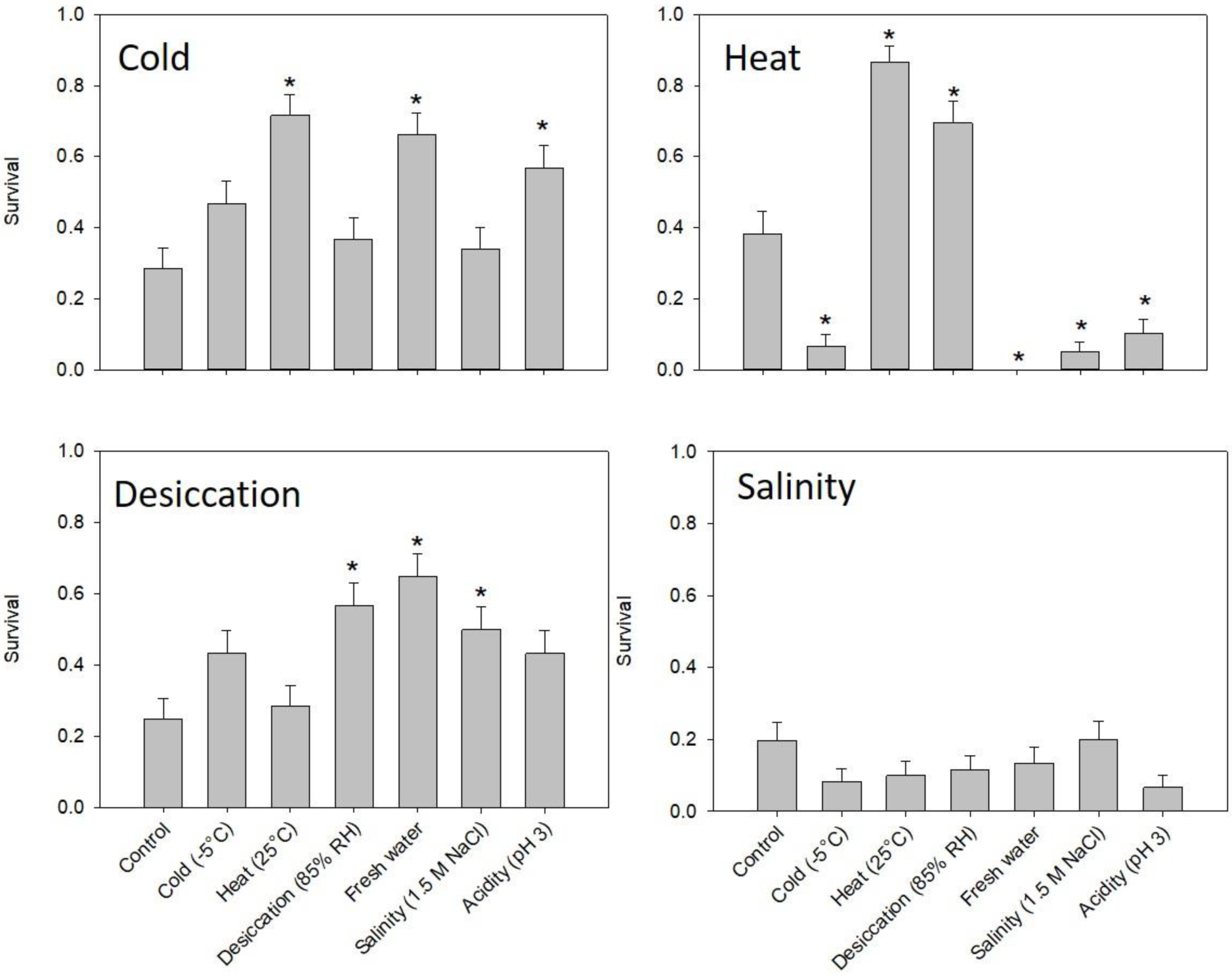
Rapid cross-tolerance in larvae of *Belgica antarctica*. Larvae were exposed to one of six 2 h pretreatments: cold (−5°C), heat (25°C), desiccation (85% RH), fresh water, salinity (1.5 M NaCl), and acidic pH (pH=3) prior to being exposed to a 24 h post-treatment of either cold (−14°C), heat (30°C), desiccation (35% RH), or salinity (4 M NaCl). Post-treatments are indicated in the top left of each panel. For each group, n=60 larvae, and asterisks indicate a significant difference between a particular group and the untreated controls (p<0.05, GLM, planned contrasts with Bonferroni correction).

## Discussion

In freezing assays, most of the treatments improved cold tolerance of larvae, with the exception starvation, heat, and changes in pH. However, of the treatments that failed to enhance organismal survival, all, with the exception of modest heat at 15°C, enhanced survival in one or both tissues, suggesting they may still offer some physiological protection against cold. While some of these stresses are known triggers of RCH (i.e. temperature and dehydration), others, such as hypoosmotic challenge, pH, UV irradiation, and starvation have not previously been linked to RCH responses. Similarly, in generalized rapid acclimation assays, many of these cues enhanced organismal survival of freezing, high temperature, or dehydration, including acidity, hypoosmotic challenge, and hyperosmotic challenge. Together, these results indicate that insects make rapid acclimatory adjustments during exposure to diverse environmental cues and that the enhanced stress tolerance resulting from these responses protect insects from other multiple abiotic stresses.

Numerous physiological and biochemical mechanisms are quickly-activated by abiotic stresses and may contribute to rapid acclimatory responses, though few of them have previously been linked specifically to RCH. For example, starvation for as few as 3 h increases autophagic proteolysis in *Drosophila melanogaster* (Scott et al., 2004), UV-B irradiation enhances total antioxidant capacity within 30 min in two species of noctuid moths (Meng et al., 2009; Karthi et al., 2014), and dehydration increased trehalose and proline synthesis in *B. antarctica* (Teets et al., 2012). Further, these mechanistic responses often confer enhanced tolerance to other abiotic stresses. In addition to starvation, autophagy may be an important part of the responses to brief chilling and dehydration (Teets and Denlinger, 2013; Gerken et al., 2015). Proline accumulation enhances freeze tolerance in insects (Koštál et al., 2011), and also increases resistance to damage from UV-B irradiation, osmotic stress, and acidosis in plants (Heuer, 1994; Kurkdjian and Guern, 1989; Alexieva et al., 2001). UV-B irradiation, dehydration, and freezing all trigger enhanced antioxidant capacity (Joanisse and Storey, 1998; França et al., 2007; Meng et al., 2009; Karthi et al., 2014). Thus, there may be mechanistic overlap among the responses triggered by disparate abiotic stresses.

This overlap in mechanisms induced by different environmental cues may help explain the cross-tolerance observed among these responses. Previous work in *B. antarctica* and many other insects demonstrates that exposure to one stress can enhance tolerance of another, which is particularly well-characterized between osmotic perturbation and low temperature (Bayley et al., 2001; Yancey, 2005; Elnitsky et al., 2009; Benoit et al., 2009; Kawarasaki et al., 2013). Yet, in this study, we detected significant effects of different rapid acclimatory responses by measuring responses to many environmental cues, where even those that were unrelated to temperature or osmotic perturbation enhanced stress. These results suggest that brief exposure to diverse environmental cues elicit generalized acclimatory responses that enhance tolerance of a multiplicity of stresses.

Interestingly, though there are similarities among rapid acclimatory responses and their underlying physiological mechanisms, recent evidence suggests that the RCH responses triggered by brief chilling (cold-induced RCH) and dehydration (drought-induced RCH) are mechanistically distinct (Yi et al., 2017). It is also possible that many other rapid acclimatory responses trigger distinct mechanisms. If so, these distinct physiological responses could presumably be induced simultaneously and may have additive effects on stress tolerance (Yi et al., 2017).

Due to their small size and correspondingly large surface area, insects are vulnerable to changing ambient conditions. Modest fluctuations in temperature can have dramatic effects on insects’ body temperature, which can, in turn, disrupt essential physiological processes (Stevenson, 1985; Heinrich, 1993). Even during brief periods of cloud cover, thoracic temperatures of foraging bumblebees can decrease enough to prevent flight (Heinrich, 1972). Further, abiotic stressors rarely occur as single, isolated pressures in nature (Holmstrup, 2010). For example, moving from shade to direct sunlight is associated with elevated UV irradiation, increased temperature, and lower relative humidity (Nicolson and Louw, 1982; Hadley, 1994), and, when microhabitats flood, temperature fluctuation and hypoxia accompany hypoosmotic challenge from direct exposure to fresh water (Hoback et al., 1998). Thus, concomitant induction of multiple rapid acclimatory responses may be ecologically important. A recent study in the flesh fly, *Sarcophaga bullata*, demonstrated that triggering both cold- and drought-induced RCH decreased the lower lethal temperature and hastened recovery of the righting reflex following cold shock (Yi et al., 2017). Because RCH has traditionally been investigated as a response to only one trigger at a time, we have likely underestimated insects’ capacity for modulating their physiological state through simultaneous induction of multiple, generalized, acclimatory responses.

In addition to naturally-occurring stresses that include freezing, dehydration, osmotic challenges, and fluctuations in pH (Lee and Denlinger, 2015), *B. antarctica* larvae are now having to overcome serious challenges from anthropogenic activity and climate change (Vaughan et al., 2003). Since the middle of the 20^th^ century, the western Antarctic Peninsula has warmed at a rate that is nearly 10 times greater than the global average, which has severe, cascading effects across trophic levels (Smith, 1994; Convey et al., 2002; Fraser and Hofmann, 2003; Smith et al., 2003; Clarke et al., 2007; Boyce et al., 2010). Warming trends are accompanied by increases in UV-B radiation, precipitation, and an elevated risk of invasion by non-native plants and animals (Convey et al., 2002; Chown et al. 2012). The Antarctic midge is small and unable to fly, therefore the potential for southward range expansion to colder regions of the Antarctic Peninsula is limited (Lee and Denlinger, 2015). Thus, tolerance of diverse abiotic stresses may be particularly important for their survival on the Antarctic Peninsula.

We used the polyextremophilic larvae of *B. antarctica* for this study because they exhibit a strong RCH response and experience a variety of stresses in nature (Lee and Denlinger, 2015). We surmised that they were likely able to make rapid acclimatory adjustments to these cues, and our results support this conclusion. Now, the answers to some critical follow-up questions are needed to assess the ecological relevance of these remarkable responses on a broader scale. Namely, are these rapid acclimatory responses 1) highly conserved throughout insect taxa, and 2) generalized in the protection they provide against other abiotic stresses? Cold-induced RCH is a highly conserved response that is found in diverse arthropod taxa (Lee and Denlinger, 2010). Its nearly-ubiquitous presence in terrestrial invertebrate ectotherms suggests that it is ecologically important to the success of these groups, which is supported by accumulating evidence in the literature (Kelty and Lee, 1999; 2001; Shreve et al., 2004; Kelty, 2007; Rajamohan and Sinclair, 2009). Other types of rapid acclimatory responses, however, are less well known, and examining their prevalence will provide clues about their ecological relevance. Exploring the question of generality will improve understanding of the extent of cross tolerance among diverse stresses and may reveal unknown or little-appreciated effects of these responses. For example, since brief heating enhances resistance to insecticides in two species of mosquitoes (Patil et al., 1996), might other types of rapid acclimation have similar effects?

The results from this study support a growing body of evidence that RCH is both triggered by numerous abiotic cues and produces a multiplicity of physiological responses, which can be altogether independent of cold (Coulson and Bale, 1991; Patil et al. 1996; Yoder, et al., 2006).

